# *In situ* NMR reveals a pH sensor motif in an outer membrane protein that drives bacterial vesicle production

**DOI:** 10.1101/2025.01.21.634179

**Authors:** Nicholas A Wood, Tata Gopinath, Kyungsoo Shin, Francesca M. Marassi

**Author notes:** **Corresponding Author:** Francesca M. Marassi, Department of Biophysics, Medical College of Wisconsin, 8701 Watertown Plank Road, Milwaukee, WI 53226.

## Abstract

The outer membrane vesicles (OMVs) produced by diderm bacteria have important roles in cell envelope homeostasis, secretion, interbacterial communication, and pathogenesis. The facultative intracellular pathogen *Salmonella enterica* Typhimurium (STm) activates OMV biogenesis inside the acidic vacuoles of host cells by upregulating the expression of the outer membrane (OM) protein PagC, one of the most robustly activated genes in a host environment. Here, we used solid-state nuclear magnetic resonance (NMR) and electron microscopy (EM), with native bacterial OMVs, to demonstrate that three histidines, essential for the OMV biogenic function of PagC, constitute a key pH-sensing motif. The NMR spectra of PagC in OMVs show that they become protonated around pH 6, and His protonation is associated with specific perturbations of select regions of PagC. The use of bacterial OMVs is an essential aspect of this work enabling NMR structural studies in the context of the physiological environment. PagC expression upregulates OMV production in *E. coli*, replicating its function in STm. Moreover, the presence of PagC drives a striking aggregation of OMVs and increases bacterial cell pellicle formation at acidic pH, pointing to a potential role as an adhesin active in biofilm formation. The data provide experimental evidence for a pH-dependent mechanism of OMV biogenesis and aggregation driven by an outer membrane protein.

**Significance:** This work sheds light on the mechanism for extracellular vesicle biogenesis by Gram negative bacteria. It shows that the *Salmonella* surface protein PagC, a major driver of extracellular vesicle formation, harbors a set of pH-sensitive histidines that become protonated at acidic pH, increasing vesicle production, and promoting bacterial cell aggregation. NMR analysis of PagC in natively secreted bacterial vesicles is introduced as a new important tool for *in situ* structural analysis of bacterial membrane proteins. The results have important implications for understanding the molecular factors that drive the formation of bacterial extracellular vesicles, their functions in human infection, as well as their roles as vaccine, drug delivery and nanotechnology platforms.

---

Extracellular vesicle secretion is a fundamental membrane remodeling process shared by eukaryotic and prokaryotic cells across all kingdoms of life. In diderm bacteria, including Gram-negative species the vesicles released from the outer membrane (OM) perform many functions important for host infection, environmental adaptation and colonization, and also have important roles as vaccine, drug delivery and nanotechnology platforms (1-3). While curvature-inducing proteins are known to play prominent roles in actively driving eukaryotic membrane remodeling and vesiculation, by scaffolding or wedging (4, 5), the induction of prokaryotic membrane curvature and OMV biogenesis is thought to be a more passive process, driven by disruption of OM-peptidoglycan connections, periplasmic accumulation of fragmented and misfolded biomolecules, or incorporation of modified LPS in the outer leaflet of the OM (1-4). In this study, we present evidence that the OM protein PagC, which has been shown to increase OM vesiculation in *Salmonella* (6-8), is an active driver of OMV biogenesis.

The expression of PagC is induced by the two-component PhoPQ regulon upon exposure to acidic pH, divalent cation limitation or cationic antimicrobial peptides (9-11). Recently (6), we showed that PagC is sufficient for driving OMV production by *Salmonella enterica* serovar Typhimurium (STm) and we localized the OMV production activity to two sequences harboring three histidines (H60, H62, H102) in the second and third extracellular loops (EL2 and EL3) of its predicted eight-stranded transmembrane β-barrel structure (**Fig. 1; Fig. S1**). This locus is an attractive candidate for a pH-sensing motif because the typical acid dissociation constant (pKa∼6) of the His sidechain makes it susceptible to protonation and positive charge acquisition in the acidic intravacuolar environment encountered by STm during infection (9-11). Given that acidic pH is one of the most potent activating signals for the PhoPQ regulon, the subset of activated genes, including PagC, may be expected to respond to environmental pH. Indeed, mimicking acidic conditions by replacing His with positively charged Lys, generates a super-activated PagC mutant that over-produces OMVs compared to wild-type, while replacing His with Ala dramatically suppresses OMV production (6). Molecular dynamics (MD) simulations indicate (6) that PagC can alter its shape and dynamics, from a cylinder at neutral pH, to a wedge at acidic pH that could induce OM curvature and vesiculation, but whether the histidines of PagC are susceptible to protonation and whether PagC undergoes conformational change have remained open questions.

**Figure 1.**
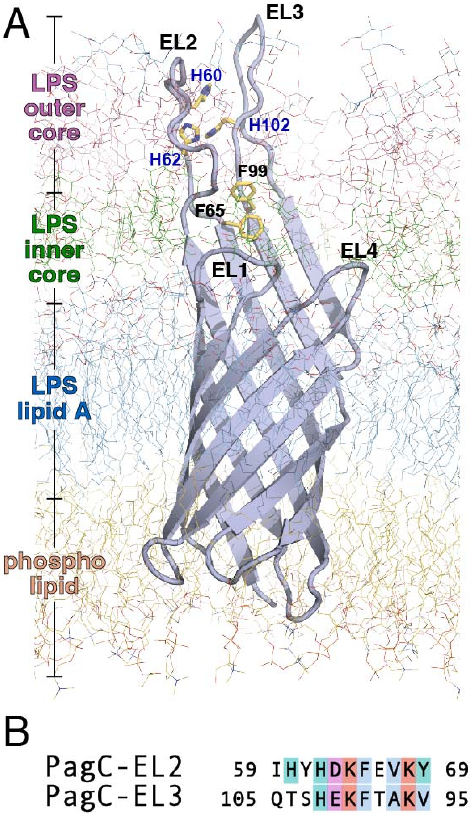
Structural model of PagC. **(A)** Representative structure of PagC, taken at 2,998 ns of MD simulation in the OM, showing the three His and two Phe side chains (yellow sticks) in the proposed pH-sensing motif. The OM includes LPS (blue, green, pink lines) and phospholipid (yellow lines). **(B)** Inverse sequence alignment of the ascending and descending neighboring segments of EL2 and EL3. Alignments are rendered with ClustalX coloring.

Here, we used solid-state nuclear magnetic resonance (NMR) and electron microscopy (EM), with native bacterial OMVs, to analyze the effects of pH on His protonation, PagC conformation and OMV structure. The data show that H60, H62 and H102 become protonated at acidic pH, and acidic pH induces specific changes in select regions of the protein. PagC expression appears to upregulate OMV production in *E. coli*, replicating its function in STm and reinforcing its role as a unique driver of OMV biogenesis. Moreover, acidic pH induces extensive OMV autoaggregation and bacterial cell pellicle formation, suggesting a potential role of PagC in biofilm formation. The data point to an active mechanism for pH-dependent modulation of OMV biogenesis by a bacterial OM protein.

## Results and Discussion

### PagC localizes to *E. coli* OMVs

Bacterial OMVs have the same membrane composition and organization as the parental bacterial OM, including an inner leaflet composed of phospholipids, an outer leaflet composed of LPS, and various OM proteins (**Fig. 1A**). This highly asymmetric membrane organization is essential for supporting protein structure and function (12, 13), yet not easily mimicked in samples reconstituted from purified components. To analyze the structure and activity of PagC in the native context, we sought to work with OMVs isolated from bacteria modified only by an inducible *pagC* gene. Since *E. coli* has been shown to support the function of PagC (14), we transformed Lemo21(DE3) *E. coli* cells with a *pagC*-encoding, IPTG-inducible plasmid, and isolated OMVs from the culture media for analysis.

SDS-PAGE and Western blots (**Fig. 2A**) show that the resulting OMVs are enriched in plasmid-encoded PagC (pPagC) relative to whole cells, as reported for STm (6). The appearance of a heat-modifiable band, that shifts from ∼14 kDa to 18 kDa upon heat denaturation, demonstrates that the protein is folded and behaves as an integral membrane β-barrel. OMVs isolated from cells transformed with empty plasmid vector (pEV), but otherwise cultured and induced as pPagC cells, do not contain PagC and therefore serve as a valuable negative control. Both pPagC and pEV OMVs are highly enriched in OmpF, as evidenced by the characteristic migration on SDS-PAGE and mass spectrometry. This trimeric porin facilitates the general passive diffusion of polar molecules (600-700 Da) across the OM (15) and is a known component of the OMV proteome (16) where it supports the exchange of solutes and protects the vesicle structures from osmotic stress.

**Figure 2.**
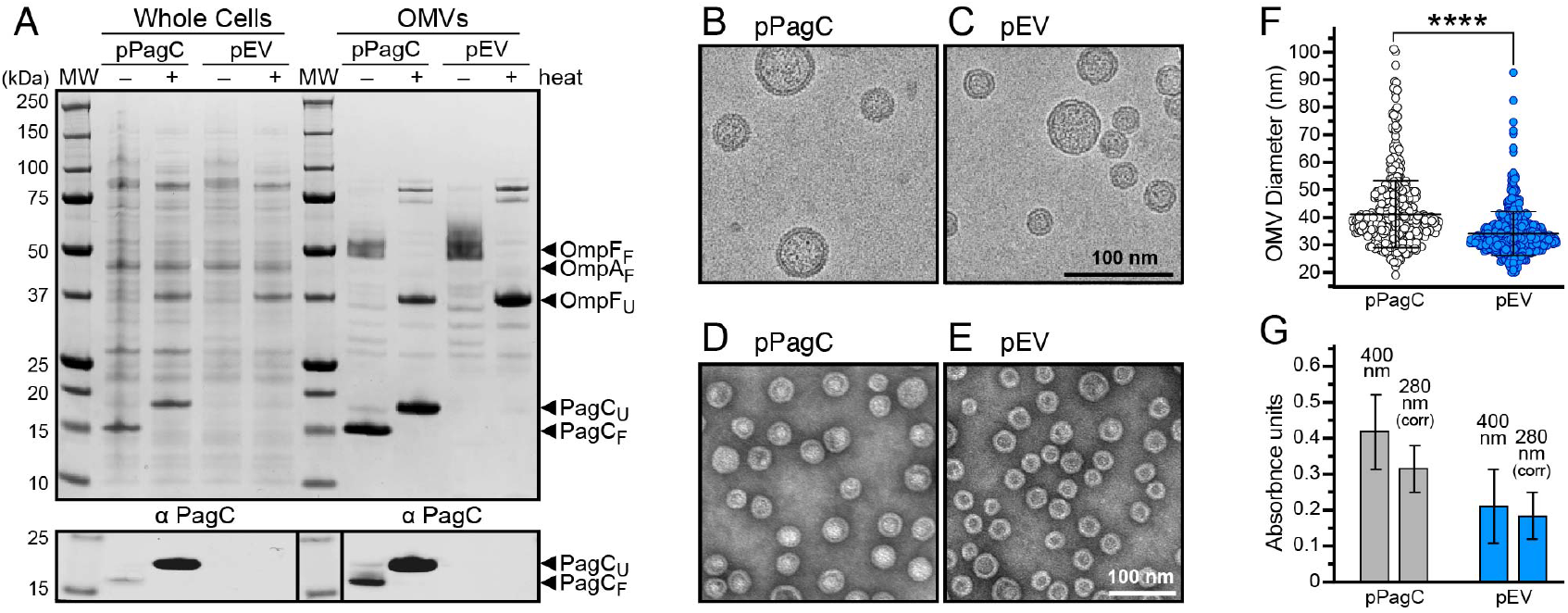
PagC expressed in *E. coli* localizes to OMVs. **(A)** SDS-PAGE of whole cells and OMVs isolated from pPagC or pEV (empty plasmid) bacteria. Proteins were visualized with Coomassie stain (top) or immunoblotting with PagC-specific antibody (α-PacC, bottom). Arrows mark bands from folded (F) and heat-unfolded (U) proteins. **(B-E)** Representative cryo-EM (B, C) or negative stain EM (D, E) of pPagC and pEV OMVs at pH 7. **(F)** OMV size estimates from negative stain EM analysis. For each of pPagC or pEV *E. coli*, 500 OMVs were selected for analysis using FIJI. Statistical significance (n≥3 biological replicates) was calculated using paired Student’s two-tailed *t* test and set at a p value <0.05 (****: p<0.0001). **(G)** Measurement of light absorbance at 400 nm and 280 nm for pPagC or pEV *E. coli*. A_280_ measurements were corrected for light scattering at 380 nm and 400 nm. Bars represent mean values with standard error of the mean error bars (7 biological replicates).

Cryogenic EM shows that both pPagC and pEV OMVs adopt discrete spherical structures (**Fig. 2B, C**), with a distinct single membrane and the two leaflets of the lipid bilayer clearly resolved as electron-dense regions, as described for OMVs from diderm bacterial (17). Negative stain EM shows that pPagC and pEV OMVs each have relatively uniform size (**Fig. 2D, E**), but PagC expression increases the average OMV diameter to 41.3 nm compared to the average of 34.2 nm observed for pEV OMVs (**Fig. 2F**). This result differs from the 165 nm diameter estimates reported for STm OMVs (8), and although we cannot exclude that our recombinant *E. coli* may yield smaller OMVs, it is also possible that differences across the two studies reflect different experimental approaches for sizing.

Nanoparticle tracking analysis, used in the STm and many other OMV studies, fails to report vesicle diameters below 60 nm (18), while an important advantage of the EM approach, used here, is the ability to select only biogenic OMVs and exclude the typically larger and often multilamellar vesicular structures that can result from explosive cell lysis (**Fig. S2**).

Finally, PagC expression appears to also result in approximately two-fold higher yield of OMVs as estimated by both light scattering and UV absorbance (**Fig. 2G**). The data therefore indicate that the OMV producing activity of PagC observed in STm (6, 8) is reproduced in *E. coli*, and provide further evidence that OMV activation is a PagC-specific property.

### Solid-state NMR analysis of PagC OMVs

Solid-state NMR is ideally suited for structural studies of integral membrane proteins in lipid assemblies, including bacterial cell envelopes and membranes (19-21). Here, we took biogenic OMVs isolated from *E. coli* directly for solid-state NMR analysis. To achieve PagC-targeted ^15^N and ^13^C isotope labeling, and eliminate NMR signals arising from endogenous *E. coli* proteins, we used a protocol where the T7/DE3-mediated expression of plasmid-encoded PagC is coupled with the suppression of chromosomal gene transcription by the antibiotic rifampin (22-24). The cells were first grown in unlabeled minimal media and only transferred to isotopically labeled media to induce protein expression. OMVs were isolated from the culture media and packed into 3.2 mm magic angle spinning (MAS) rotors for NMR experiments.

The ^15^N and ^13^C spectra of pPagC OMVs (**Fig. 3, black**), obtained with ^1^H-^15^N, ^1^H-^13^C and ^1^H-^15^N-^13^C cross-polarization (CP), contain signals from protein, as well as lipid and LPS acyl chains, glycans, aminoglycans and potentially peptidoglycan. These essential cellular building blocks continue to be produced and incorporate ^13^C and ^15^N during the isotope labeling period of cell culture and are known components of the bacterial OMV lumen (1-3). Their NMR signals have been observed in the spectra of cell envelope samples (21, 25, 26) and could be assigned by comparison.

**Figure 3.**
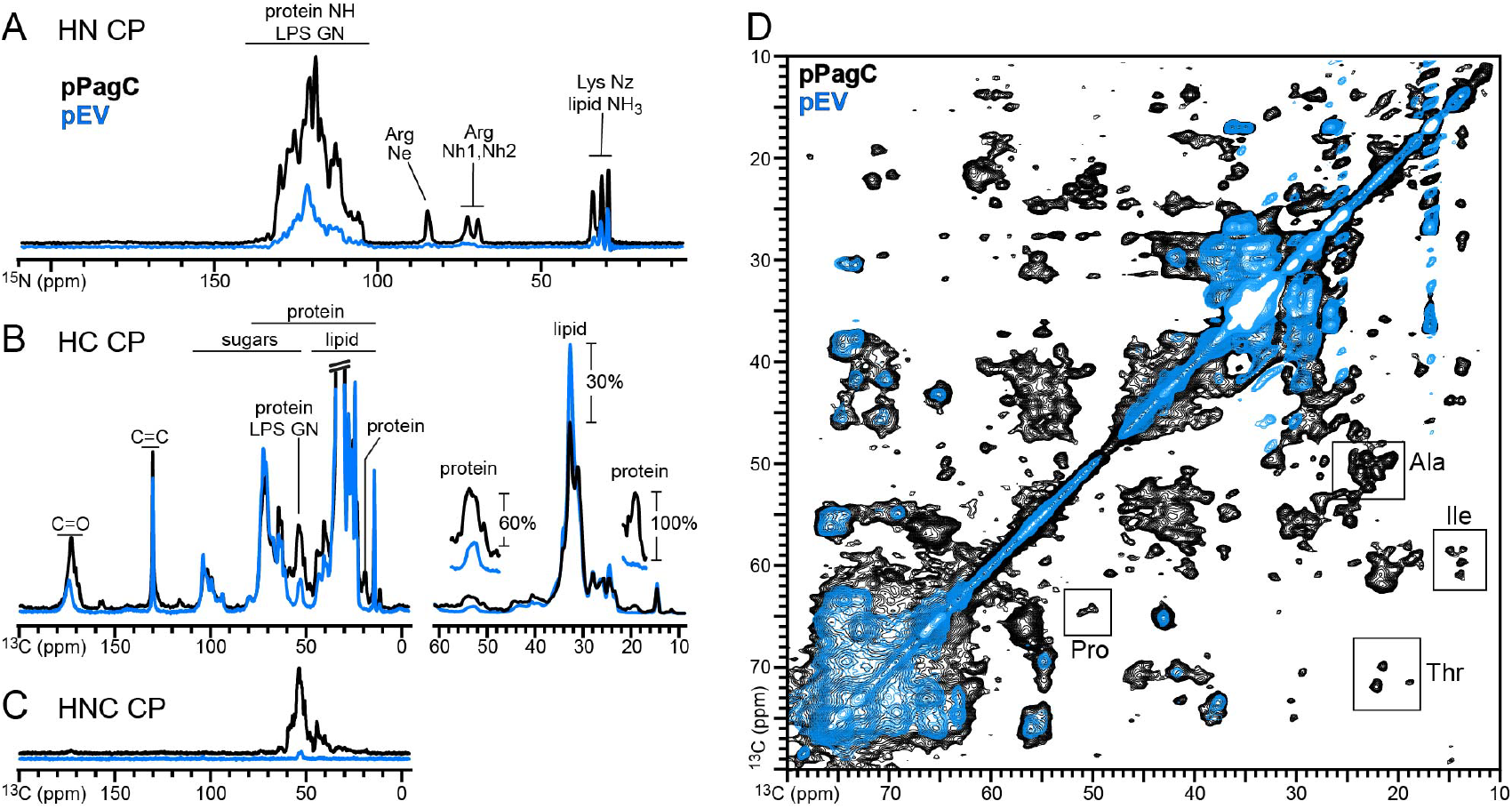
NMR spectra of PagC in *E. coli* OMVs. Spectra were acquired at 4°C for pPagC (black) or pEV (blue) OMVs at pH 7. **(A-C)** One-dimensional spectra acquired with ^1^H-^15^N CP (A), ^1^H-^13^C CP (B), or ^1^H-^15^N followed by ^15^N-^13^C CP (C). Signals from protein and LPS sites are marked (LPS GN: glucosamine). **(D)** Two-dimensional ^13^C/^13^C correlation PDSD spectra. Assignments to residue types are marked in inset boxes.

By contrast, little evidence of protein NMR signals is seen in the spectra of pEV OMVs, isolated from cells transformed with empty plasmid vector, but otherwise grown and induced in an identical manner as pPagC cells (**Fig. 3, blue**). The ^13^C spectrum obtained by ^1^H-^13^C CP has both greater lipid signal intensity at 33 ppm (∼30%), as well as much lower protein signal at 54 ppm (∼60%) and 19 ppm (100%), for pEV versus pPagC OMVs (**Fig. 3B**), and the ^13^C spectrum obtained by ^1^H-^15^N-^13^C polarization transfer shows a dramatic signal reduction **(Fig. 3C)**.

The two-dimensional ^13^C/^13^C correlation spectrum of pPagC OMVs (**Fig. 3D, black**) has many resolved signals from Ala, Ile, Pro, Ser, and Thr spin systems. These can be easily identified based on their characteristic chemical shifts at positions consistent with β-barrel structure. By contrast, the spectrum of pEV OMVs (**Fig. 3D, blue**) has no detectable signals from protein, confirming that the expression system leads to highly selective isotope labeling of the target with minimal background, notwithstanding the abundance of OmpF and other proteins in the OMVs.

### The histidines of PagC are sensitive to pH

The proposed pH sensing motif of PagC comprises two inverse homology sequences (**Fig. 1**) each located in the descending segment of EL2 (residues 62-69) and ascending segment of EL3 (residues 95-102). MD simulations (6) revealed a role for these sites: in forming a series of EL2-EL3 cross contacts, including H60/H62-H102 and F65-F99 ring stacking, that stabilize the loop conformations and couple their dynamics, while at low pH, electrostatic repulsions from protonated His decouple EL2 and EL3, altering both the conformation of PagC and its interactions with OM lipids and LPS. This model aligns with the known low pH activation of PagC and the His mutagenesis data (6), but neither His protonation in the bacterial OM, nor the existence of an EL2-EL3 pH sensing motif that can switch conformation upon transfer to acidic environments, has been explored experimentally.

To address the question of His protonation, we acquired ^15^N NMR spectra for a series of pPagC OMV samples incubated in buffers ranging from pH 8.5 to pH 4 (**Fig. 4A**). The ^15^N chemical shift tensors of His imidazole Nd and Ne nitrogens are well characterized, and isotropic ^15^N chemical shifts are good indicators of both His protonation and hydrogen bonding (27-30).

**Figure 4.**
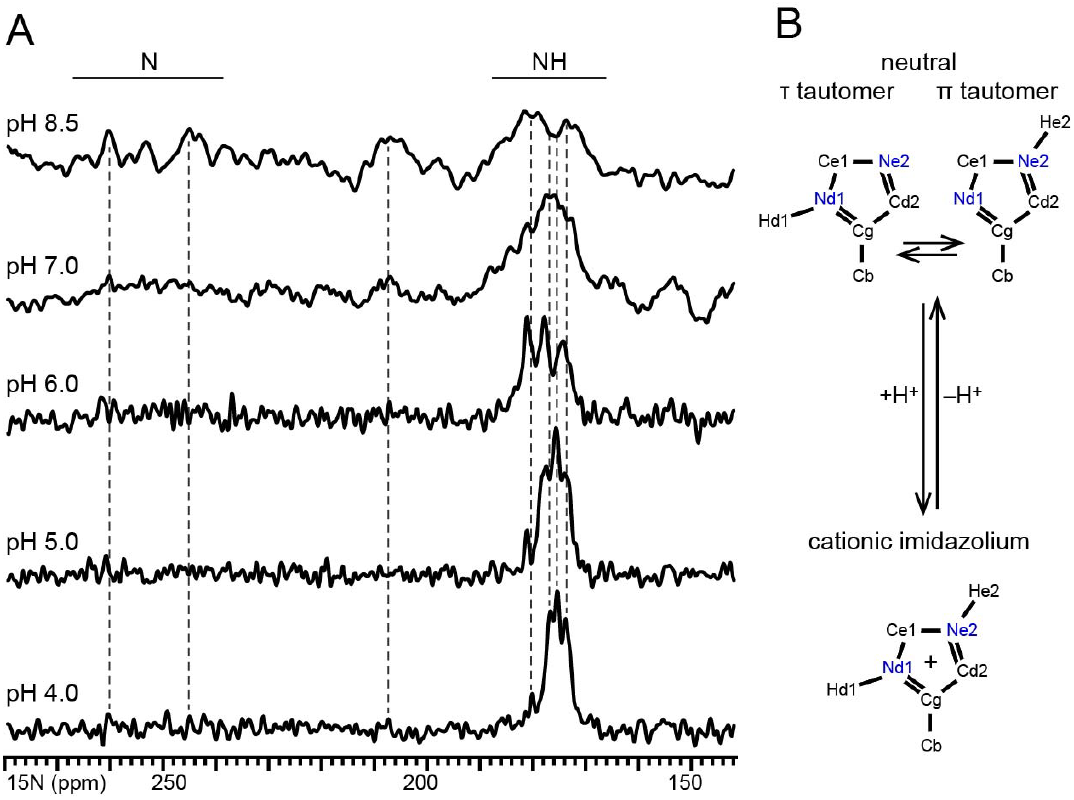
pH Titration of PagC His. **(A)** One-dimensional ^15^N spectra of pPagC OMVs acquired at 4°C with ^1^H-^15^N CP. Dashed lines align with resolved signals. **(D)** Structures of the tautomeric states of the neutral His imidazole ring and fully protonated imidazolium cation.

Environmental pH has a dramatic effect on the ^15^N spectra of the His imidazole nitrogens of PagC. At pH 8.5, a broad envelope of signal intensity is observed between 172 and 260 ppm. Based on reported values of His ^15^N chemical shifts (27-30), the 172-182 ppm intensity may be assigned to protonated Ne and Nd, while the weak 245-260 ppm signal likely reflects deprotonated nitrogens which are more challenging to detect by ^1^H-^15^N CP due to the lack of bound hydrogen. Overall, the absence of resolved peaks, combined with the weak broad signal intensity near 207 ppm, reflect a mix of neutral tautomeric states, with either Nd (τ tautomer) or Ne (π tautomer) protonation, exchanging slower than the μs timescale of the ^15^N NMR experiment (**Fig. 4B**).

As the pH is lowered, the 207 ppm intensity disappears, and resolved spectral components between ∼172-181 ppm grow in intensity with progressive line narrowing. Three signals resolved at pH 6 and four resolved at pH 5 and pH 4, may be assigned to protonated Ne and Nd of the imidazolium cation. The signal observed at 180.1 (pH 4) and 181.1 ppm (pH 6 and pH 5) is consistent with hydrogen bonding of a protonated imidazole nitrogen to a carboxylate group (30).

The spectra are complex. The observation of three to four ^15^N signals for three His may reflect rapid imidazole ring rotations that average the two chemically inequivalent Ne and Nd sites of each His, spectral overlap of distinct Ne and Nd signals, or selective detection of specific His residues with favorable hydrogen bonding or dynamic characteristics for NMR observation by CP. Absent resonance assignment, it is impossible to distinguish among these mechanisms. Nevertheless, the data unequivocally demonstrate that the His of PagC are water-accessible and sensitive to environmental pH and protonation near pH 6.

### Effect of pH on the conformation of PagC

To probe the effect of pH on other protein sites, we acquired ^15^N/^13^C and ^13^C/^13^C correlation spectra of pPagC OMVs exposed to either pH 7 or pH 4 buffer (**Fig 5A, B**). Both ^15^N/^13^C and ^13^C/^13^C spectra compare favorably with those of the PagC homolog Ail in the bacterial cell envelope (21, 31), demonstrating the potential of NMR characterization of membrane proteins in biogenic OMVs. The NMR signal intensity is ∼ 50% greater at pH 4 than pH 7, and the number of scans were adjusted to compensate for this difference (Table S1), which likely arises from a strong OMV aggregation effect (see below) observed at pH 4 allowing us to pack more sample into the rotor, and some enhancement of ^1^H-^13^C CP due to increased protonation of exchangeable sites.

**Figure 5.**
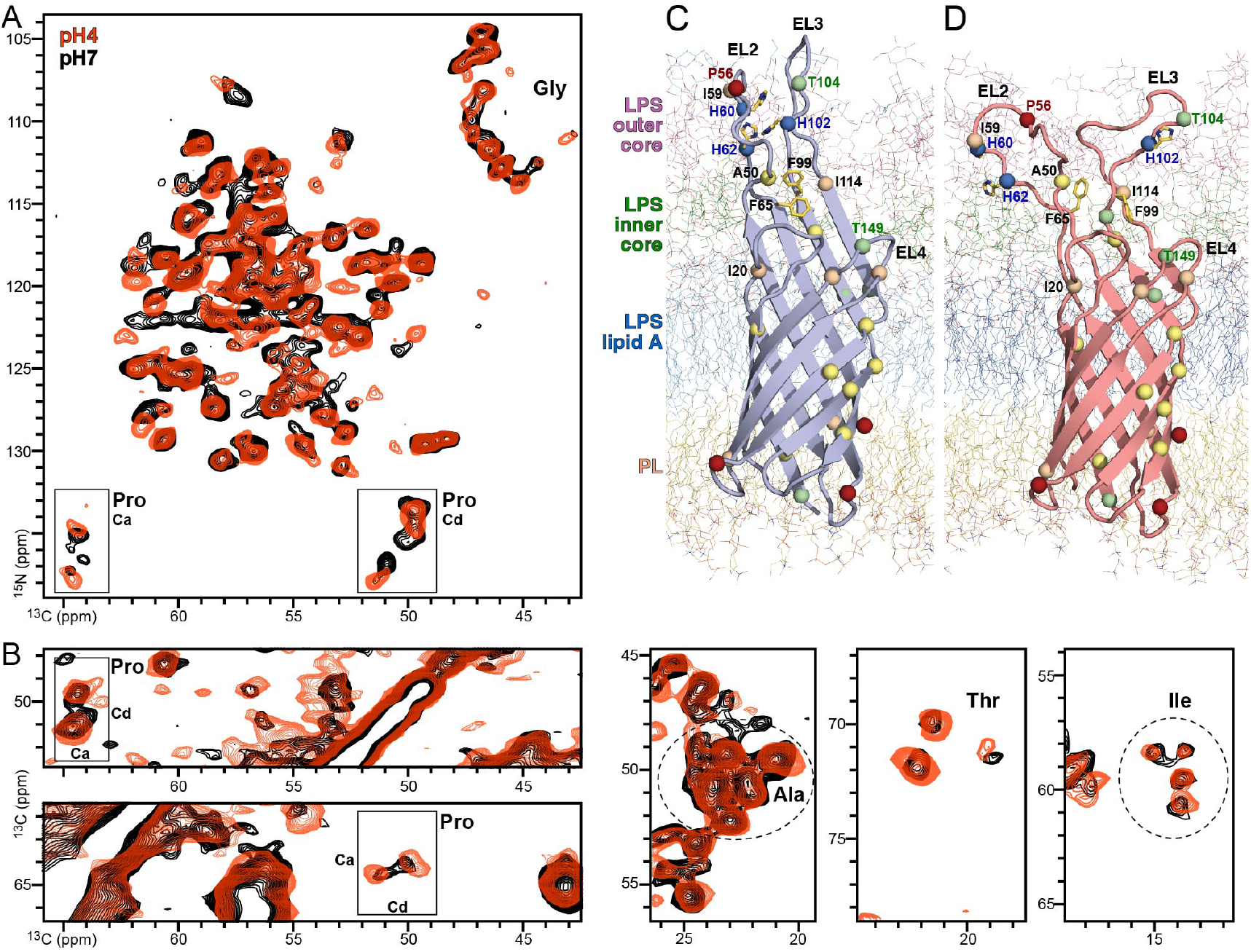
Effect of pH on the ^15^N/^13^C and ^13^C/^13^C NMR spectra of PagC in *E. coli* OMVs. Spectra were acquired at 4°C for pPagC OMVs at pH 7 (black) or pH 4 (red). **(A)** Two-dimensional ^15^N/^13^C TEDOR NCA spectra. **(B)** Two-dimensional ^13^C/^13^C PDSD spectral regions selected to show signals from Pro, Ala, Thr and Ile. **(C, D)** Structural models taken from MD simulations of His-neutral (C; 2,998 ns) or His-protonated (D; 2,668 ns) PagC, showing His and Phe sidechains (yellow stick), and CA atoms (sphere) of Ala (yellow), Ile (wheat), Pro (red), and Thr (green). The OM includes LPS (blue, green, pink lines) and phospholipid (PL: yellow lines).

The ^15^N/^13^C spectra acquired with transfer echo double resonance (TEDOR) mixing have excellent signal intensity and resolution, with line widths of well-resolved signals in the range of 0.6-0.8 ppm for ^13^C, and 1.3-1.8 ppm for ^15^N (**Fig. S3**). Many signals can be resolved from the eighteen Gly, four Pro, six Thr, nine Ala and seven Ile in the sequence, including Pro N-Ca and N-Cd correlations. Since these residues are distributed across the length of the PagC β-barrel their NMR signals are valuable probes of protein conformation (**Fig. 5C, D**).

The ^15^N and ^13^C chemical shifts from these sites reflect β-stranded secondary structure, and the spectra obtained at pH 7 and pH 4 display extensive signal overlap, indicating that the overall conformation of PagC is maintained irrespective of pH. Notably, there are also several ^15^N and ^13^C chemical shift perturbations of select signals. For example, two Thr (T98, T104) are located in EL3 and two Thr signals are perturbed by pH in the ^13^C/^13^C spectrum (**Fig. 5B**). Moreover, select signals from hydrophobic Ala and Ile, and at least two of the Pro Ca and Cd signals, are perturbed in response to pH. While the lack of resonance assignments precludes a detailed spectral analysis, such selective peak perturbation reflects a site-specific response to the local environment of neighboring protein and LPS moieties, rather than a wholesale response to buffer pH. Whether this response includes changes in conformation will require structure determination.

### PagC induces OMV autoaggregation and pellicle formation at acidic pH

In the process of NMR sample preparation, we noticed a marked flocculation of pPagC OMVs occurring immediately upon transfer to pH 4, and this prompted us to examine the ultrastructural impact of pH. Negative stain EM of freshly prepared OMVs reveals striking aggregation of pPagC OMVs, but not pEV OMVs, at acidic pH (**Fig. 6A**). While all OMVs are well-dispersed at pH 7.5, pPagC OMVs lightly cluster at pH 6, and the dominant features at pH 5 and pH 4 are extensive aggregates, with clear evidence of membrane fusion and some structures resembling wire-like OMV connections (**Fig. 6A, arrow**) similar to those described for *Myxococcus xanthus* (32). While we could identify some solitary non-aggregated vesicles, their abundance was substantially reduced relative to neutral pH. Aggregation appears to initiate around pH 6 and complete fusion is observed at pH 4.

**Figure 6.**
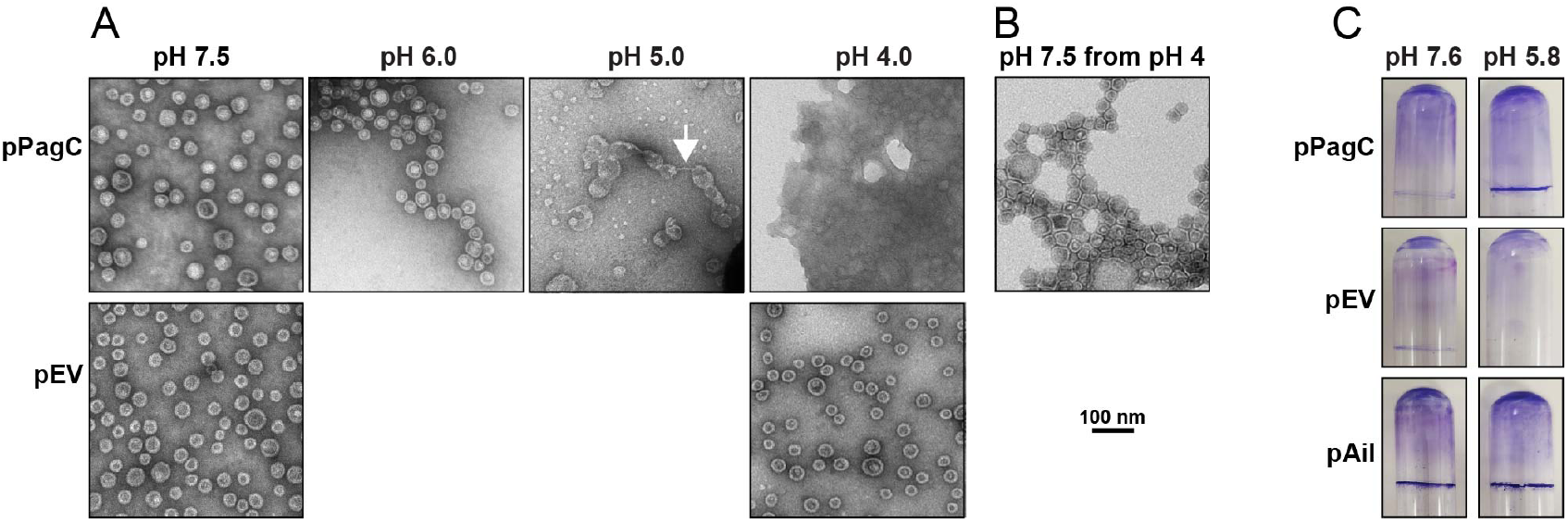
PagC induces pH-dependent OMV aggregation and bacterial cell pellicle formation. **(A, B)** Representative negative stain EM of pPagC (top) and pEV (bottom) OMVs, acquired at pH 7.5, pH 6, pH 5, pH 4 (A) or pH 7.5 after incubation at pH 4 (B). White arrow points to a wire-like OMV connection at pH5. **(C)** Pellicle formation by pPagC, pEV and pAil *E. coli* cells. Cells were suspended in 2 mL of M9 minimal media (OD_600_=0.5) in 15×100 mm glass tubes, and incubated for 16 h at 37°C. After removing the cells by centrifugation, the interior glass walls were treated with methanol and air dried overnight, then washed three times with buffer, treated with 2.5 mL of crystal violet solution (0.1% for 10 min), then washed with buffer and air dried. Pellicle formation was detected as a violet-stained rim at the air-water interface.

To test the reversibility of this phenomenon, we performed EM on pPagC OMVs incubated first at pH 4 overnight and then switched to pH 7.5 for another 12 hours (**Fig. 6B**). While many aggregate masses remain, some are visibly dissolved, suggesting some degree of reversibility. Tightly associated OMVs appear to form a mosaic-like structure in which the vesicle membranes flatten against each other increasing the contact surface. Moreover, some membranes appear to have fused, suggesting not only that PagC promotes aggregation but also that it may mediate membrane fusion.

We also tested the ability of PagC to induce *E. coli* cell pellicle formation at pH 7.6 and pH 5.8 where its OMV producing activity in STm was first tested (6). At pH 7.6 there is minimal evidence of pellicle formation at the air-water interface, but a substantial pellicle forms at pH 5.8 as readily visualized by staining with crystal violet (**Fig. 6C; Fig. S6**). By contrast, pEV cells form very little pellicle at either neutral or acidic pH, while pAil cells produce dense pellicles regardless of pH, in line with the documented aggregation activity of Ail (21, 33).

These results align with the reported association of PagC with biofilm formation and multiple drug resistance in *Salmonella enterica* (34), as well as growing evidence that OMVs are common constituents of the microbial biofilm extracellular matrix and play an active role in biofilm formation (35-39). The pH-dependent reversible aggregation may reflect a mechanism for environmental adaptation whereby secreted OMVs aggregate to form an adhesive matrix at low pH, that can be quickly undone upon transfer to a neutral pH environment. Such a process could be important for the survival of *Salmonella* in environments of extreme pH fluctuation, such as the human digestive tract.

The aggregation phenotypes remain to be demonstrated in STm. Non-typhoidal *Salmonella* produce an alternative group IV O-antigen capsule that is implicated in environmental survival, immune evasion, and biofilm formation (40-43), while the LPS core of *E. coli* B strains, including Lemo21(DE3) used in this study, lacks O-antigen repeats and capsular biosynthesis (44, 45) and thus, may artificially unmask PagC exposing its extracellular loops to the OMV surface and activating aggregation.

### PagC is an active driver of OMV biogenesis

OMV secretion is a major membrane remodeling process essential for bacterial colony behavior and environmental adaptation. Recently (6), we proposed a model for the pH-dependent OMV biogenic activity of PagC where His protonation at acidic pH drives conformational change from an ordered cylindrical barrel to a wedge, where EL2 and EL3 are decoupled and exert pressure on the outer leaflet of the OM resulting in curvature and vesicle budding. Here, we have shown that the histidines of PagC are indeed pH-sensitive and that their protonation is associated with perturbations of specific PagC sites. The solid-state NMR spectra show that PagC adopts an overall β-stranded conformation in bacterial OMVs, and the His sidechains titrate from neutral tautomeric states to the protonated imidazolium cation near pH 6. Whether Asp and Glu residues in the extracellular loops are also susceptible to protonation and contribute to protein conformational change is an interesting question that needs to be addressed in future studies. Overall, the NMR data reflect a specific response of PagC to environmental pH and provide experimental support for an active role as a pH sensor and driver of OM curvature.

The use of bacterial OMVs is an essential aspect of our study enabling structural analysis in the context of the native environment. Working with native samples is particularly important for bacterial OM proteins, which have co-evolved with LPS to confer species-specific properties, and function as a coordinated assembly with the highly asymmetric environment of OM (12, 13). Here, the OMV producing capacity of PagC observed in STm was recapitulated in a B strain *E. coli* derivative. Most interestingly, the propensity of PagC to promote OMV aggregation and bacterial pellicle formation at low pH is an important new property of this protein. With the caveat that the R1 type LPS of our B strain *E. coli* may unmask the PagC aggregation phenotype, this finding aligns with evidence for the association of PagC with biofilm formation and multiple drug resistance in *Salmonella enterica* (34)

OMVs isolated from engineered *E. coli* have been proposed as an *in situ* platform for solution NMR studies of periplasmic proteins or periplasmic domains of membrane proteins (46). This application, however, relied on the deletion of multiple OM protein genes and vesicle size homogenization by filtered extrusion. Moreover, the use of solution NMR excludes the possibility of examining integral membrane proteins whose dynamics are coupled to the relatively large OMV structure. Here, we took native OMVs, isolated from *E. coli* with the full complement of endogenous, chromosomally encoded OM proteins, directly for solid-state NMR analysis. This work, therefore, paves the way for NMR studies of physiological bacterial OMVs including those produced by STm.

In conclusion, our work provides experimental evidence for the role of PagC as an active pH sensor and driver of OMV biogenesis, describes a potential new role of PagC as a regulator of OMV aggregation and pellicle formation, and introduces OMVs as an important new platform for *in situ* NMR structural analysis of OM proteins.

## Materials and Methods

### Bacterial strains, culture conditions, and key reagents

All *E. coli* were commercially available Lemo21(DE3) (New England BioLabs, MA, USA). The pET22b(+)-*pagC* plasmid (pPagC) for IPTG-inducible expression of untagged PagC lacking the endogenous signal peptide was designed in frame with the PelB signal peptide (47) and acquired from Genscript (Piscataway, NJ, USA). Empty vector bacteria were transformed with empty pET22b(+) plasmid (pEV). Western blots were performed with a custom (Thermo Fisher Scientific, IL, USA) polyclonal antibody against the PagC EL3 peptide CDGDSFSNKISSRKTGFAWG. Bacteria were cultured in M9 media supplemented with 0.2% glucose, 100 µg/mL ampicillin and 35 µg/mL chloramphenicol when appropriate.

### Isotope labeling of PagC

For isotopic labeling, transformed bacteria were grown overnight at 37°C in unlabeled M9 media. Overnight cultures were used to inoculate fresh unlabeled M9 to achieve OD_600_=0.1, and cultures were incubated with shaking at 37°C. At OD_600_=0.6, expression of the DE3 RNA polymerase was induced by adding 0.4 mM IPTG. Following a 10-minute incubation, the cells were pelleted (5,000xg, 4°C, 20 min), and resuspended in M9 supplemented with 1 g/L of ^15^N ammonium sulfate and 0.2% ^13^C glucose (Cambridge Isotope Laboratories, MA, USA). For selective labeling, IPTG and rifampin were added to 0.4 mM and 100 µg/mL, respectively. Expression was allowed to proceed for 16 hours at 26° C prior to harvesting of samples. Empty vector control samples were cultured identically to PagC-containing samples.

### OMV isolation and characterization

OMVs were harvested by diafiltration and ultracentrifugation. Briefly, all cultures were centrifuged to remove bacteria, and the resulting supernatants were centrifuged again to remove any residual bacteria. This supernatant was filtered through a 0.45 µm filter to remove residual bacteria and large aggregates. Filtrates were concentrated using an Amicon stirred cell and an appropriate 300 kDa PES membrane disc filter (Millipore Sigma, St. Louis, MO, USA). Concentrated filtrates were then ultracentrifuged at 150,000 xg, 4° C for 1 hour and 30 minutes, after which the OMV pellets were resuspended in the desired buffer and centrifuged again. OMV pellets were then resuspended in the desired buffer and used for subsequent analysis. All buffers for OMV resuspension contained 0.9 mM CaCl_2_ and 0.5 mM MgCl_2_ to preserve OMV membrane integrity as previously described (48). OMVs were quantified by light scattering and UV absorbance, as described (48, 49).

### Cryo-EM

Purified OMVs (3 *μ*L of ∼1 mg/mL) were applied to a glow-discharged Quantifoil 300-mesh gold grid in a Vitrobot chamber (ThermoFisher Mark IV). Excess liquid was blotted with a blot force of 3, blot time of 6 s, and the grid was plunge-frozen into liquid ethane. Imaging was performed on a ThermoFisher Glacios 200 kV electron microscope equipped with a Falcon 4 direct electron detector, and Selectris energy filter. Micrographs were collected with a magnification of 100 kx, a pixel size of 1.089 Å/pixel, and a total dose of 60 e-/Å^2^. Images were processed using Cryosparc v4.4.

### Negative stain EM

Samples were adsorbed onto glow-discharged 400 mesh Formvar carbon copper grids (Electron Microscopy Sciences, PA, USA) for 15 sec, and then wicked away with filter paper. The grids were then treated with 2% uranyl-acetate for 30 sec, then blotted to remove the stain, and allowed to dry for at least one hour prior imaging at 100,000x magnification on a Jeol JEM-1400 Transmission Electron Microscope (JEOL USA, MA) at 60 kV. For each experiment, 500 OMVs were selected form measurement using FIJI (50).

### Solid-state NMR spectroscopy

OMV samples were centrifuged into 3.2 mm zirconium rotors for Solid state NMR. Solid state NMR experiments were performed on a 700 MHz NeO Bruker spectrometer, equipped with a 3.2 mm E-free ^1^H/^13^C/^15^N probe. All experiments were acquired at a sample temperature of 4°C, with a MAS rate of 12.5 kHz, and a recycle delay of 2 s. The 90° pulse lengths for ^1^H, ^13^C, and ^15^N were set to 3 µs, 6 µs and 6 µs. Heteronuclear decoupling was achieved using the spinal-64 pulse sequence with 70 kHz radiofrequency amplitude. During ^1^H-^13^C and ^1^H-^15^N CP, 30 kHz radiofrequency amplitude was used on ^13^C or ^15^N channels. The Hartmann-Hahn matching condition was set by ramping the ^1^H radiofrequency amplitude to maximize signal intensity. During double ^1^H-^15^N-^13^C CP, NCA polarization transfer was obtained with 22 kHz and 32 kHz radiofrequency spinlocks on the ^13^C and ^15^N channels, with 70 kHz continuous wave decoupling on the ^1^H channel. For the TEDOR experiments, Hadamard encoding (51) was applied with 2 ms selective I-BURP pulses on the ^15^N channel. NMR experimental details are provided in Table S1.

### Pellicle formation

Pellicle formation was assayed as described (21) with minor modifications. Briefly, cells from an overnight induction pelleted, washed with sterile PBS, and resuspended in non-buffered M9. OD600 was then recorded, and the volume of cell suspension necessary to achieve an OD600 of 0.5 in 2 mL was pelleted and resuspended in M9 media buffered to either pH 7.6 or pH 5.8. These cultures were transferred to 15 × 100 mm glass tubes and were incubated at 37° C with gentle agitation for 16 h. For supplementation with either pEV or pPagC OMVs, 100 µg (determined by adjusted A_280_ calculations) of purified OMVs were added during resuspension of the cells prior to incubation. Following incubation, cell suspensions were aspirated. Then, 2.5 mL of 100% ice cold methanol was added to each tube, incubated for 5 minutes, and removed. Tubes were air dried overnight. The next day, tubes were washed three times with PBS prior to staining for 10 minutes with 0.1% crystal violet. After removal of the crystal violet solution, stained pellicles were rinsed twice with PBS to remove excess dye. Tubes were air dried overnight prior to imaging. Pellicle formation was determined qualitatively as a crystal violet stained ring at the air-liquid interface.

## Acknowledgments

This work was supported by grants from the National Institutes of Health (GM118186, AI188770 and F32 AI188770). It utilized NMR instrumentation supported by a grant from the NIH (OD028716), transmission EM instrumentation supported by the MCW-Oxford Instruments Center for Advanced Microscopy and Electron Microscopy Core, and cryo-EM instrumentation supported by the MCW Cancer Center. We thank Ashish Gadicherla and Linda Olson for their assistance with EM, and Marassi lab members for discussion.

## Author contributions

NAW, TG, KS and FMM were involved in research design, data analysis and discussion. NAW and TG performed experiments. NAW, TG, KS and FMM wrote the manuscript with input and approval from all authors.

## Supplementary Information

**Table 1.**
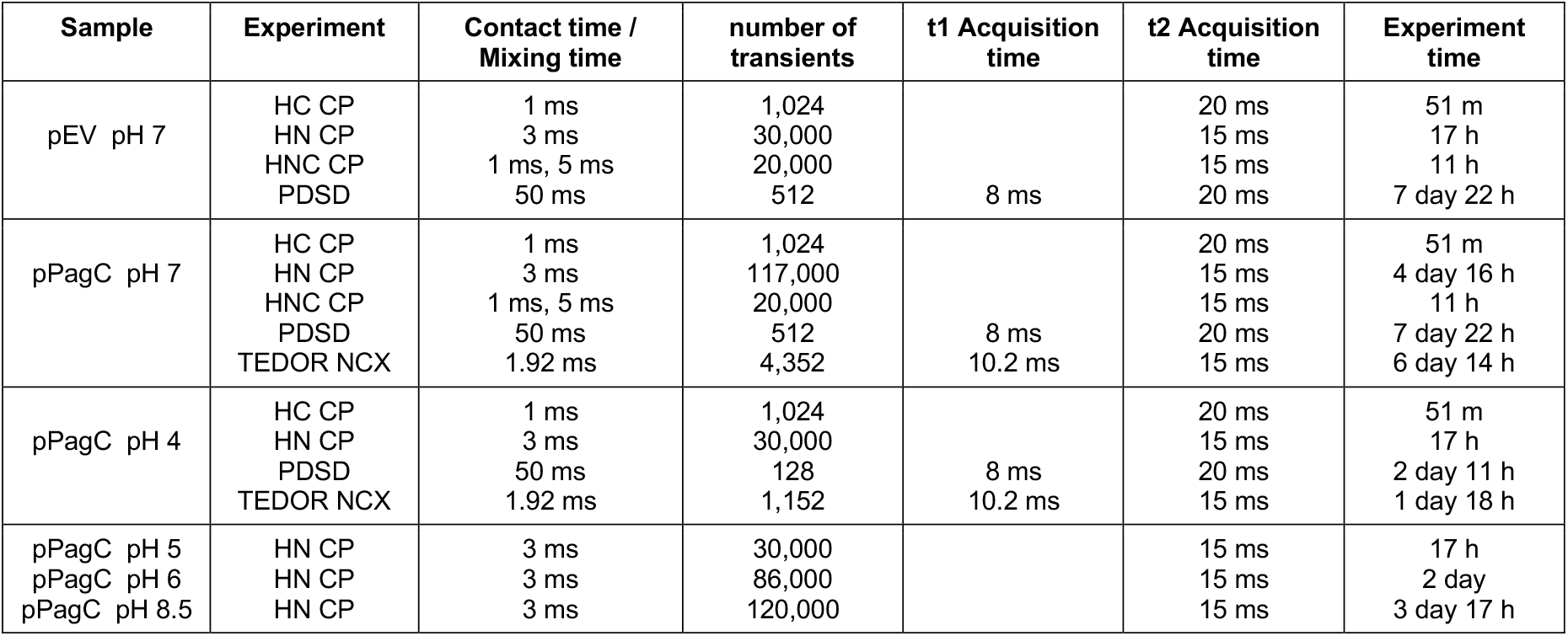
NMR Samples and solid-state NMR experimental parameters.

**Figure S1.**
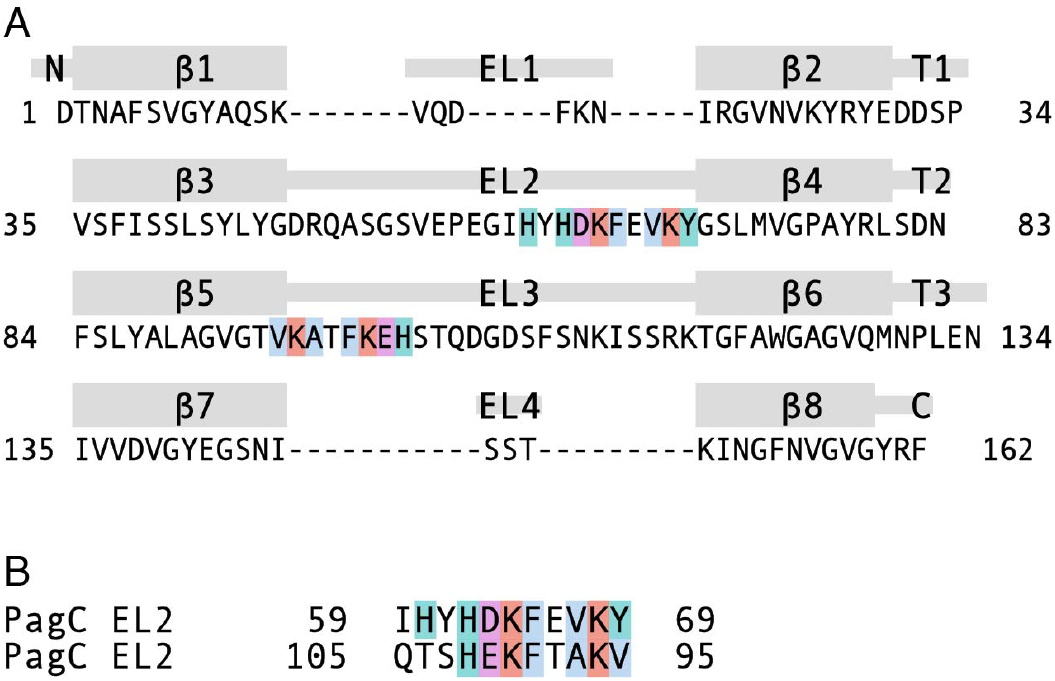
Structure-aligned sequence of PagC. **(A)** Sequence of PagC aligned to the predicted β strands, extracellular loops (EL1-EL4) and intracellular turns (T1-T4). Neighboring segments of EL2 and EL3 that harbor the His are colored. **(B)** Inverse sequence alignment of the ascending and descending neighboring segments of EL2 and EL3. Alignments are rendered with ClustalX coloring.

**Figure S2.**
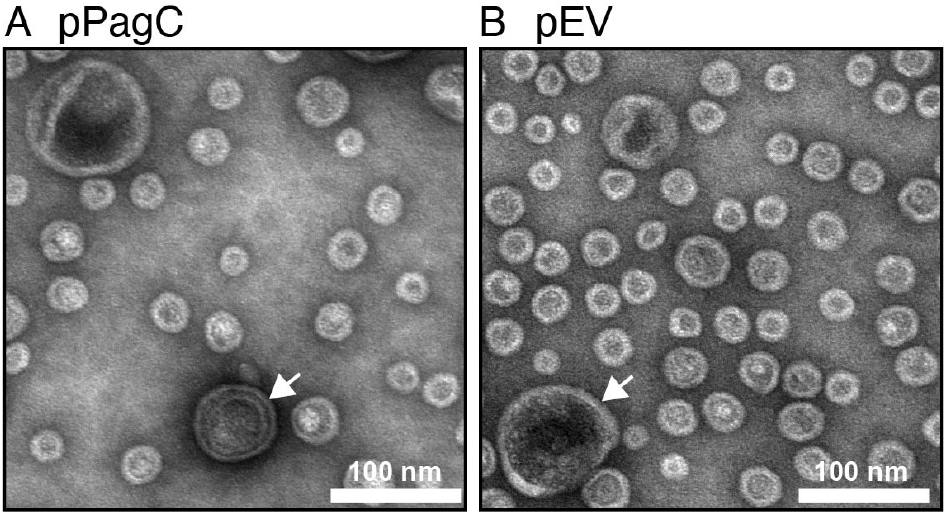
Representative negative stain TEM of OMVs at pH7. **(A)** pPagC OMVs. **(B)** pEV OMVs. Arrows denote OMVs with double or multiple membranes (A), or enclosing numerous smaller vesicles within the lumen (B). These were excluded from size analysis.

**Figure S3.**
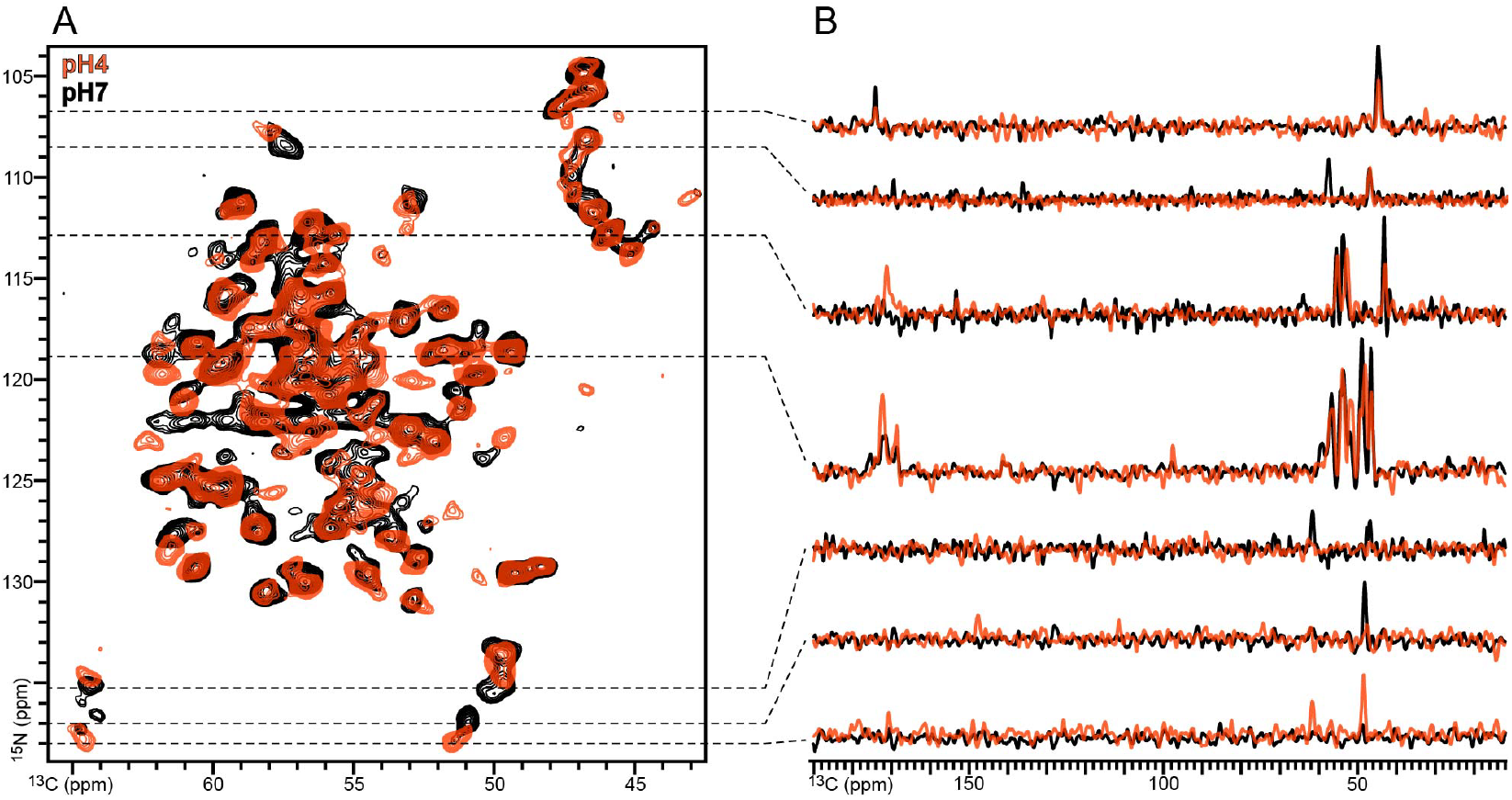
Effect of pH on the ^15^N/^13^C NMR spectrum of PagC in *E. coli* OMVs. Spectra were acquired at 4°C for pPagC OMVs at pH 7 (black) or pH 4 (red). **(A)** Two-dimensional ^15^N/^13^C TEDOR NCA spectra. **(B)** One-dimensional spectra taken from the 2D spectrum at ^15^N chemical shifts marked by the dashed lines.

**Figure S4.**
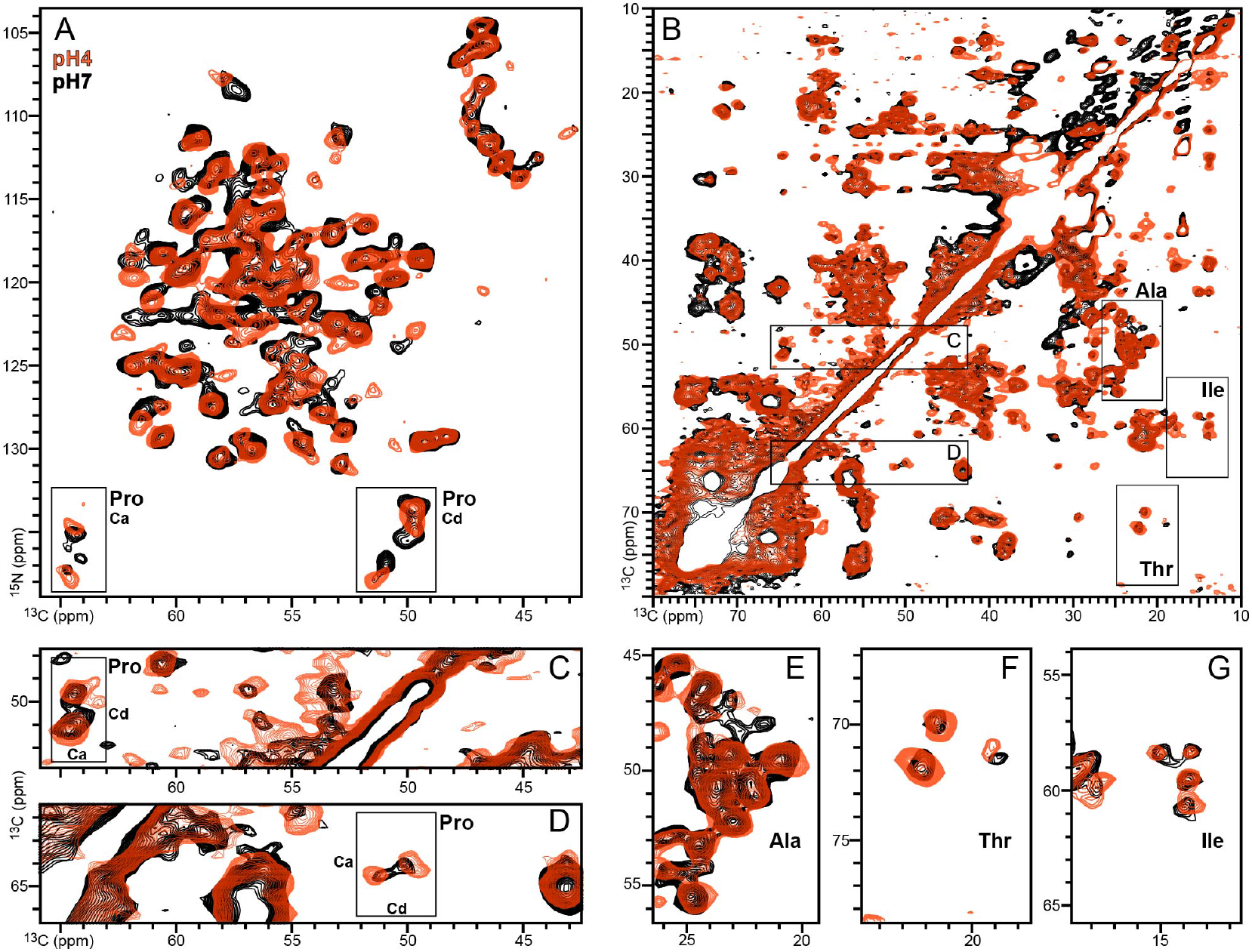
Effect of pH on the ^15^N/^13^C and ^13^C/^13^C NMR spectra of PagC in *E. coli* OMVs. Spectra were acquired at 4°C, for pPagC OMVs at pH 7 (black) or pH 4 (red). **(A)** Two-dimensional ^15^N/^13^C TEDOR NCA spectra. **(B)** Two-dimensional ^13^C/^13^C PDSD spectra. Boxes outline selected for signals from Pro, Ala, Thr and Ile. **(C-G)** Expanded regions of the ^13^C/^13^C correlation spectrum showing signals for Pro, Ala, Thr and Ile.

**Figure S5.**
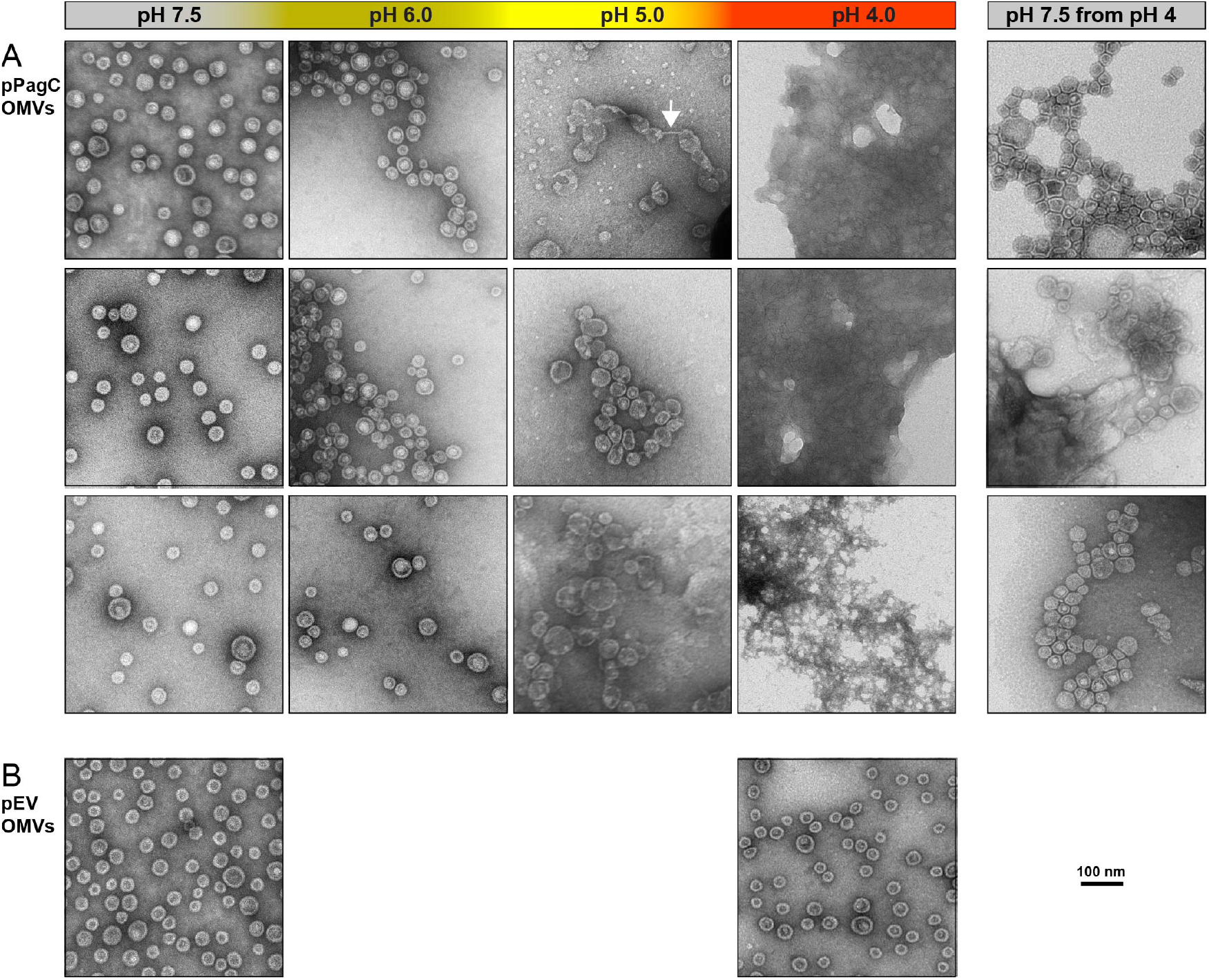
PagC induces pH-dependent OMV aggregation. **(A, B)** Representative negative stain EM of pPagC (A) and pEV (B) OMVs, acquired at pH 7.5; pH 6; pH 5; pH 4; or pH 7.5 after incubation at pH 4. White arrow points to a wire-like OMV connection at pH5. **(B)** Pellicle formation by pPagC, pEV and pAil *E. coli* cells. Cells were suspended in 2 mL of M9 minimal media (OD_600_=0.5) in 15×100 mm glass tubes, and incubated for 16 h at 37°C. After removing the cells by centrifugation, the interior glass walls were treated with methanol and air dried overnight, then washed three times with buffer, treated with 2.5 mL of crystal violet solution (0.1% for 10 min), then washed with buffer and air dried. Pellicle formation was detected as a violet-stained rim at the air-water interface.

**Figure S6.**
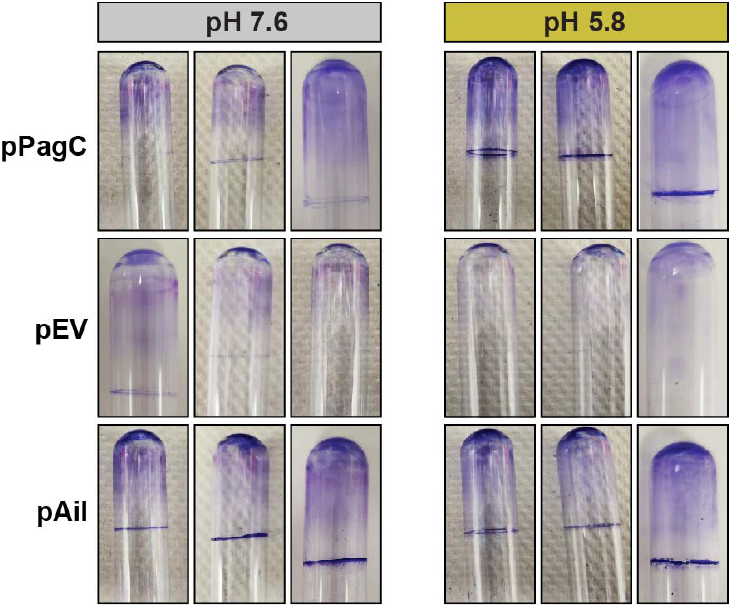
PagC induces pH-dependent bacterial cell pellicle formation. Cells (3 biological replicates) were suspended in 2 mL of M9 minimal media (OD_600_=0.5) in 15×100 mm glass tubes, and incubated for 16 h at 37°C. After removing the cells by centrifugation, the interior glass walls were treated with methanol and air dried overnight, then washed three times with buffer, treated with 2.5 mL of crystal violet solution (0.1% for 10 min), then washed with buffer and air dried. Pellicle formation was detected as a violet-stained rim at the air-water interface.

